# Adolescent onset of volitional ethanol intake normalizes sex differences observed with adult-onset ethanol intake and negative affective behaviors during protracted forced abstinence

**DOI:** 10.1101/2025.03.06.641859

**Authors:** Caitlyn M. Edwards, Zishuo Xu, Danny G. Winder

## Abstract

**Rationale:** Negative affect during ethanol abstinence can lead to relapse and dependence. Voluntary ethanol drinking models are crucial for examining negative affect following chronic ethanol access, but female rodents often drink more ethanol than males, complicating comparisons between sexes. Since chronic adolescent ethanol use poses a substantial risk for later alcohol use disorder, we hypothesize that adolescence is a critical window for consolidating drinking behavior before major hormonal changes affect ethanol consumption.

**Objectives:** This study compared sex differences in voluntary ethanol consumption and negative affective behavior in mice that initiated ethanol consumption during early adolescence (∼PND30) or early adulthood (>PND49).

**Methods:** Male and female C57BL/6J mice underwent the Chronic Drinking Forced Abstinence (CDFA) paradigm, with the “Ethanol” group given two-bottle choice access to ethanol and water, and the “Water” group given two water bottles. Ethanol intake and preference were measured over six weeks. Two weeks following ethanol removal, mice underwent behavioral testing for negative affective-like behavior.

**Results:** Adult-onset female mice consumed significantly more ethanol and displayed higher ethanol preference compared to adult-onset male mice. In contrast, adolescent-onset male and female mice consumed similar ethanol levels and displayed similar preference. We observed increased immobility during the forced swim test in adult-onset ethanol females, but not males, during protracted abstinence. However, both sexes of adolescent-onset ethanol mice displayed increased immobility during forced abstinence.

**Conclusions:** These findings highlight adolescence as a critical period during which both sexes voluntarily consume ethanol and are equally vulnerable to the behavioral disturbances associated with ethanol abstinence.

## Introduction

Adolescence is a particularly vulnerable period of brain development in which substance use is thought to produce enduring effects (Squeglia et al. 2009; Lees et al. 2020). Research indicates that early initiation of ethanol use is associated with a range of negative outcomes, including the development of alcohol use disorder (AUD) (Grant and Dawson 1997; DeWit et al. 2000). Particularly, individuals who first begin using ethanol in early adolescence (between ages 11 and 14) are more likely to become dependent on ethanol later in life (DeWit et al. 2000). Given the importance of adolescence as a critical developmental window, studying voluntary ethanol consumption initiated during this time frame and its impact on behavior has significant clinical relevance.

Negative emotional states are prevalent among individuals with AUD, particularly during periods of withdrawal and abstinence (Koob and Volkow 2016). Anxiety and depression experienced during abstinence are major drivers of relapse where individuals seek ethanol to alleviate negative affect. Negative reinforcement as a driver of ethanol consumption is particularly associated with drinking patterns in women. Women report more mentally unhealthy days following binge drinking compared to men (Wen et al. 2012) and women with AUD tend to have more comorbid psychiatric disorders (Conway et al. 2006; Greenfield et al. 2010; Hilderbrand and Lasek 2018).

The development of negative affect-like behavior during abstinence in females has also been observed with preclinical models of voluntary ethanol consumption (Pang et al. 2013; Holleran et al. 2016; Vranjkovic et al. 2018; Neira et al. 2022). These observations in females could indicate an underlying biological sex difference in proclivity to the development of behavioral disturbances during alcohol abstinence. However, there are also distinct sex differences in patterns of ethanol consumption. In most models of voluntary ethanol consumption, females of many strains of rats and mice tend to drink significantly more than males (Yoneyama et al. 2008; Centanni et al. 2018; Fulenwider et al. 2019; Dao et al. 2020; Bloch et al. 2020; Arnold et al. 2023; Perry et al. 2025; Yoon et al. 2025). Therefore, the levels of ethanol consumption could explain the observed differences in negative affect-like behavior.

Sex-related differences in ethanol consumption are likely related to differences in sex hormones (Lund and Jacobsen 1990; Satta et al. 2018). Given that prepubertal adolescents have not yet reached sexually mature sex hormone levels (Vigil et al. 2016) and that initiation of ethanol use during this time is a critical window for development of AUD (DeWit et al. 2000), we sought to examine whether initiation of voluntary ethanol consumption during early adolescence would reduce sex differences in consumption observed during adulthood. We then examined sex differences in negative affective behavior during forced abstinence in animals that initiated voluntary drinking during early adolescence and early adulthood. By comparing the effects of adolescent-onset and adult-onset drinking, this study aims to elucidate the nuanced ways in which early ethanol exposure influences mental health outcomes across sexes. This comparison will provide valuable insights into the mechanisms underlying ethanol-related negative affect.

## Methods

### Animals

Female and male C57BL/6J mice were bred in house and were given voluntary access to ethanol beginning in either early adolescence (PND30±3; n = 45 [22 females; 23 males]) or early adulthood (>PND49; n = 47 [23 females; 24 males]). All mice were single housed at the start of the Chronic Drinking Forced Abstinence (CDFA) paradigm and given *ad libitum* access to standard rodent chow (5L0D PicoLab Laboratory Rodent Diet, LabDiet). All experiments were conducted in accordance with the National Institutes of Health Guide for the Care and Use of Laboratory Animals and were reviewed and approved by the Vanderbilt University Institutional Animal Care and Use Committee.

### Chronic Drinking Forced Abstinence (CDFA) Paradigm

To measure voluntary drinking, mice were singly housed with continuous access to two 50mL conical tubes fitted with sippers described previously (Holleran et al. 2016; Centanni et al. 2018; Taylor et al. 2024). After habituating to the bottle set up for 5 days with water in both bottles, half of the mice (“Ethanol” groups) were given two-bottle choice between ethanol and water whereas the control mice (“Water” groups) continued to receive water in both bottles. Ethanol access began with an ethanol ramp: 3 days of 3% ethanol followed by 7 days of 7% ethanol. At the end of the ramp, mice had constant access to 10% ethanol for four weeks.

Bottle weights were measured every 48-72 hours to measure ethanol intake and ethanol preference over the six-week continuous ethanol access period. Bottle placement was switched at the time of bottle weight measurements to account for individual side bias. Empty cages with ethanol and water bottles were used to estimate leakage due to evaporation or movement of the cage rack. Ethanol intake was quantified as grams of ethanol consumed relative to body weight per day (g/kg/day). Ethanol preference was calculated based on the amount of ethanol consumed relative to total amount of fluid consumed from both bottles (ethanol and water). For the purposes of calculating bottle preference, one water bottle for mice in the water only group was arbitrarily assigned as the “ethanol” bottle.

Following the six-weeks of chronic drinking, mice were placed into “forced abstinence” where the ethanol bottle was removed so that all groups had access only to water.

### Behavior Tests

Two weeks following removal of ethanol, mice underwent a series of daily affective behavior tests to examine changes in affective state during forced abstinence.

#### Light/Dark Box

On day 14 of forced abstinence, mice were placed into the light side of a light/dark box (11 x 11.5 in light side; 11 x 6.5in dark side) and allowed to freely explore for 15 minutes. Behavior was recorded with a video camera. Latency to enter the dark box and time spent in the light box were quantified using the automated AnyMaze software and compared between groups.

#### Elevated Plus Maze

On day 15 of forced abstinence, mice were placed on the open arm of an elevated plus maze consisting of two open arms and two enclosed arms (each 12 x 2.5in) extending from a central platform (2.5 x 2.5in). Mice were allowed to freely explore the maze for 10 minutes while behavior was recorded with a video camera. Latency to exit the open arm and time spent on the open arms were quantified using the automated AnyMaze software and compared between groups.

#### Forced Swim Test

On day 16 of forced abstinence, mice were placed into a beaker of water (23-25°C) for 6 minutes while their behavior was recorded with a video camera. Latency to first immobility and time spent immobile were quantified manually from recorded video files using Behavioral Observation Research Interactive Software (BORIS). Immobility was defined as the complete absence of all movement. Time immobile data from one mouse (adolescent-onset ethanol male) was excluded due to a lighting malfunction during the test.

### Statistical Analysis

Data were analyzed using GraphPad Prism 10.3. Ethanol intake was analyzed in ethanol access groups using a repeated measures two-way ANOVA with Sex and Time as independent variables. During the final week of 10% ethanol access, an unpaired t-test was used to examine differences between sexes. Ethanol preference was analyzed using a repeated measures three-way ANOVA with Sex, Ethanol Group, and Time as independent variables. During the final week of 10% ethanol access, ethanol preference was analyzed using a two-way ANOVA with Sex and Ethanol Group as independent variables. For ethanol drinking behavior data, Tukey’s posthoc comparisons were used to compare individual groups. Negative affective behaviors in the light/dark box (latency to enter dark box and time in light box), the elevated plus maze (latency to exit open arm and time in open arms), and the forced swim test (latency to immobility and time immobile) were quantified using two-way ANOVAs with Sex and Ethanol Group as independent variables. For affective behavior data, Fisher’s Least Significant Difference tests were used to compare individual groups. Estimation statistics were used to report effect sizes for both drinking behavior and negative affective behavior during forced abstinence (Calin-Jageman and Cumming 2019; Ho et al. 2019).

## Results

### Voluntary Ethanol Consumption During Continuous Ethanol Access

#### Adult-Onset Ethanol Intake

First, we sought to replicate previous observations of sex differences in ethanol intake in mice that began voluntary ethanol consumption in adulthood. We compared ethanol intake relative to body weight across the CDFA paradigm between sexes in mice that began drinking in early adulthood. Repeated measures two-way ANOVA (Sex x Time) indicated a significant main effect of Time [F(5,110) = 73.52; p < 0.0001], a significant main effect of Sex [F(1,22) = 14.84; p = 0.0009], and a significant interaction between Time and Sex [F(5,110) = 3.938; p = 0.0025] on ethanol intake in adult mice (Fig. 1A). Tukey’s multiple comparisons tests indicated a significant increase in ethanol consumption in females compared to males across all 10% ethanol weeks [10% Week 1: p < 0.0001; 10% Week 2: p = 0.0006; 10% Week 3: p = 0.0016; 10% Week 4: p < 0.0001]. Unpaired t-test of the final 10% ethanol week indicated a significant difference between male and female ethanol consumption [t(22) = 5.005; p < 0.0001] (Fig. 1B). Examining the unpaired mean difference between male and female mice using estimation statistics indicated that adult female mice drank 5.86 g/kg/day more than adult male mice [95%CI 3.63, 7.92].

**Fig. 1.**
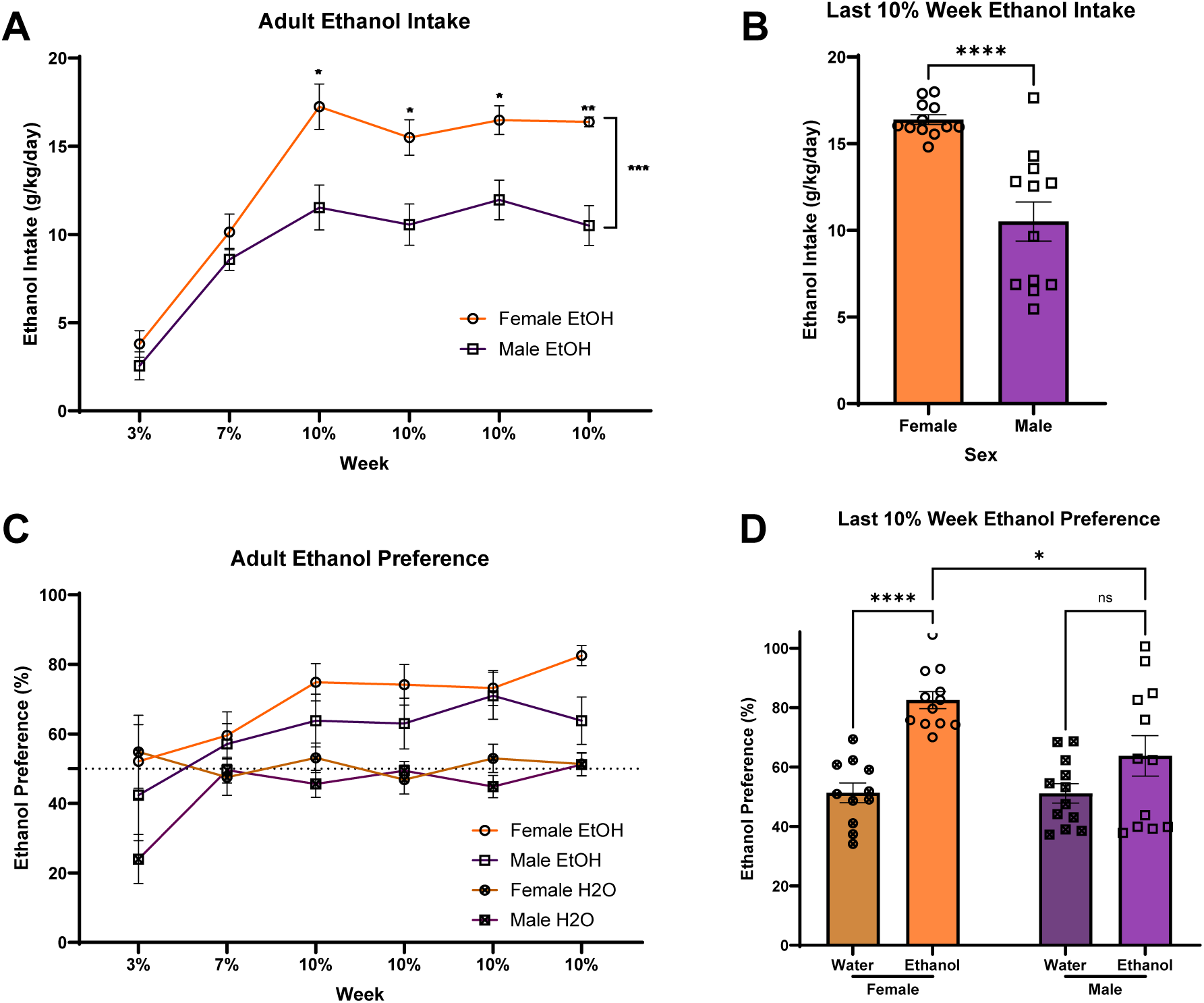
Adult-onset female mice drink consistently more ethanol and have stronger preferences for ethanol than adult-onset male mice. A) Ethanol intake relative to body weight across the CDFA paradigm. Females consumed significantly more ethanol starting from the first week of 10% ethanol compared to males. B) In the final week of CDFA, female mice drank an average of 5.88g/kg/day more ethanol than male mice. C) Preference for the “ethanol” bottle relative to the water bottle across the CDFA paradigm. Females develop a stronger preference for ethanol compared to males. D) In the final week of CDFA, female ethanol mice develop a strong ethanol preference compared to water only controls, whereas male ethanol mice do not. ^ns^p>0.05; *p<0.05; **p<0.01; ***p<0.001; ****p<0.0001.

#### Adult-Onset Ethanol Preference

We also compared preference for the ethanol bottle over the water bottle across the CDFA paradigm between sexes in mice that began drinking in early adulthood. Repeated measures three-way ANOVA (Sex x Ethanol Group x Time) indicated a significant main effect of Time [F(5,215) = 6.425; p < 0.0001], a significant main effect of Ethanol Group [F(1,43) = 18.52; p < 0.0001], and a significant main effect of Sex [F(1,43) = 4.106; p = 0.0490] on adult ethanol bottle preference (Fig. 1C). Two-way ANOVA (Sex x Ethanol Group) indicated a significant main effect of Sex [F(1,43) = 4.474; p = 0.0402], a significant main effect of Ethanol Group [F(1,43) = 24.36; p < 0.0001], and a significant interaction between Sex and Ethanol Group [F(1,43) = 4.676; p = 0.0362] on ethanol bottle preference during the final week of CDFA (Fig. 1D). Tukey’s multiple comparisons test indicated a significant increase in “ethanol” bottle preference in the ethanol group compared to water controls in females (p < 0.0001), but not males (p = 0.2103), as well as a significantly greater preference for ethanol in the females compared to the males (p = 0.0192).

#### Adolescent-Onset Ethanol Intake

We next sought to determine whether these sex differences in ethanol intake across the CDFA paradigm also exist in mice that began drinking during early adolescence. Repeated measures two-way ANOVA (Sex x Time) indicated a significant main effect of Time [F(5,100) = 71.06; p < 0.0001] but no main effect of Sex or interaction between Time and Sex on ethanol intake in adolescent mice (Fig. 2A). We directly compared ethanol intake of adolescent-onset mice and adult-onset mice. Two-way ANOVA (Sex x Age) indicated a significant main effect of Sex [F(1,42) = 22.54; p < 0.0001], a significant main effect of Age [F(1,42) = 8.274; p = 0.0063], and an interaction between Sex and Age [F(1,42) = 4.464; p = 0.0406] on ethanol intake during the final week of CDFA (Fig. 2B). Tukey’s multiple comparisons tests indicated significantly elevated ethanol intake in adult-onset ethanol females compared to males (p < 0.0001), but no difference in ethanol intake between adolescent-onset females and males (p = 0.2768), and significantly greater ethanol intake in males that began CDFA during adolescence compared to those who began during adulthood (p = 0.0055).

**Fig. 2.**
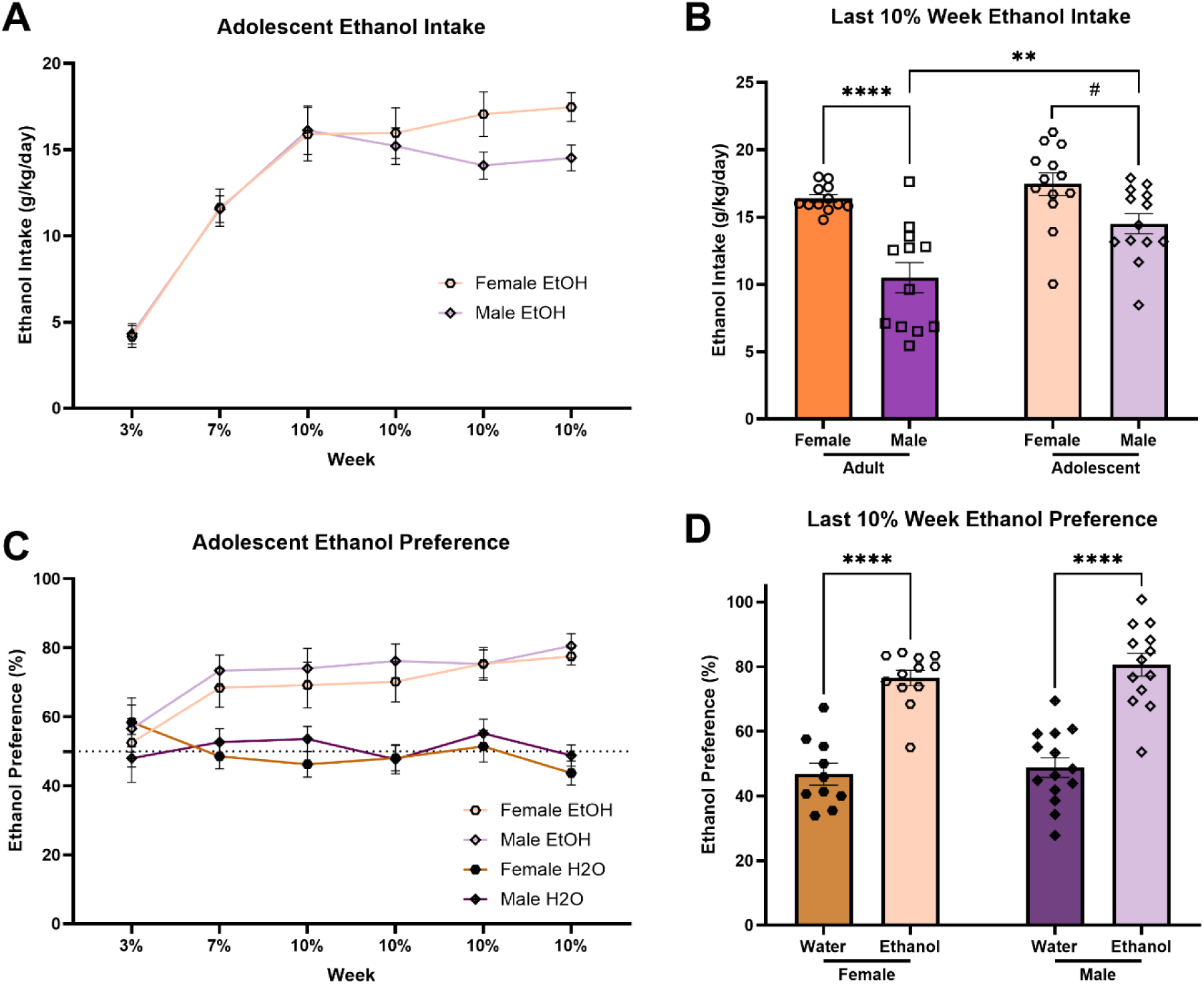
Mice that begin CDFA during early adolescence show similar levels of ethanol intake and preference between sexes. A) Ethanol intake relative to body weight across the CDFA paradigm. B) In the final week of CDFA, adolescent male mice drank significantly more ethanol compared to adult male mice while drinking similar levels to adolescent female mice. C) There are no significant sex differences in the development of preference for the ethanol bottle. D) Both female and male mice that began drinking during early adolescence demonstrated significant preference for ethanol in the final week of CDFA. ^#^p<0.10; **p<0.01; ****p<0.0001.

#### Adolescent-Onset Ethanol Preference

We also examined whether male and female adolescent-onset mice display similar preference for ethanol across the CDFA paradigm. Repeated measures three-way ANOVA (Sex x Ethanol Group x Time) indicated a significant main effect of Time [F(5,205) = 3.610; p = 0.0038], a significant main effect of Ethanol Group [F(1,41) = 46.00; p < 0.0001], and a significant interaction between Time and Ethanol Group [F(5, 205) = 7.728; p < 0.0001] on ethanol bottle preference (Fig. 2C). There was no significant main effect of Sex. Two-way ANOVA (Sex x Ethanol Group) indicated a significant main effect of Ethanol Group [F(1,41) = 89.52; p < 0.0001], but no main effect of Sex or interaction between Sex and Ethanol Group on adolescent-onset ethanol bottle preference during the final week of CDFA (Fig. 2D). Tukey’s multiple comparisons test indicated a significant increase in “ethanol” bottle preference in the ethanol group compared to water controls in both females (p < 0.0001) and males (p < 0.0001) with no significant differences between sexes.

### Negative Affective-Like Behavior During Protracted Forced Abstinence

#### Negative Affective-Like Behaviors Following Adult-Onset Ethanol

Two weeks following removal of the ethanol bottle, we tested mice for ethanol abstinence-induced negative affect-like behaviors. We observed no differences in behaviors in the light/dark box or elevated plus maze of adult-onset mice. Two-way ANOVAs (Sex x Ethanol Group) indicated no significant differences in either latency to enter the dark box and time in the light box in the light/dark box (Fig. 3A,B) and no significant differences in either latency to exit the open arm and time in the open arms in the elevated plus maze (Fig. 3C,D).

**Fig. 3.**
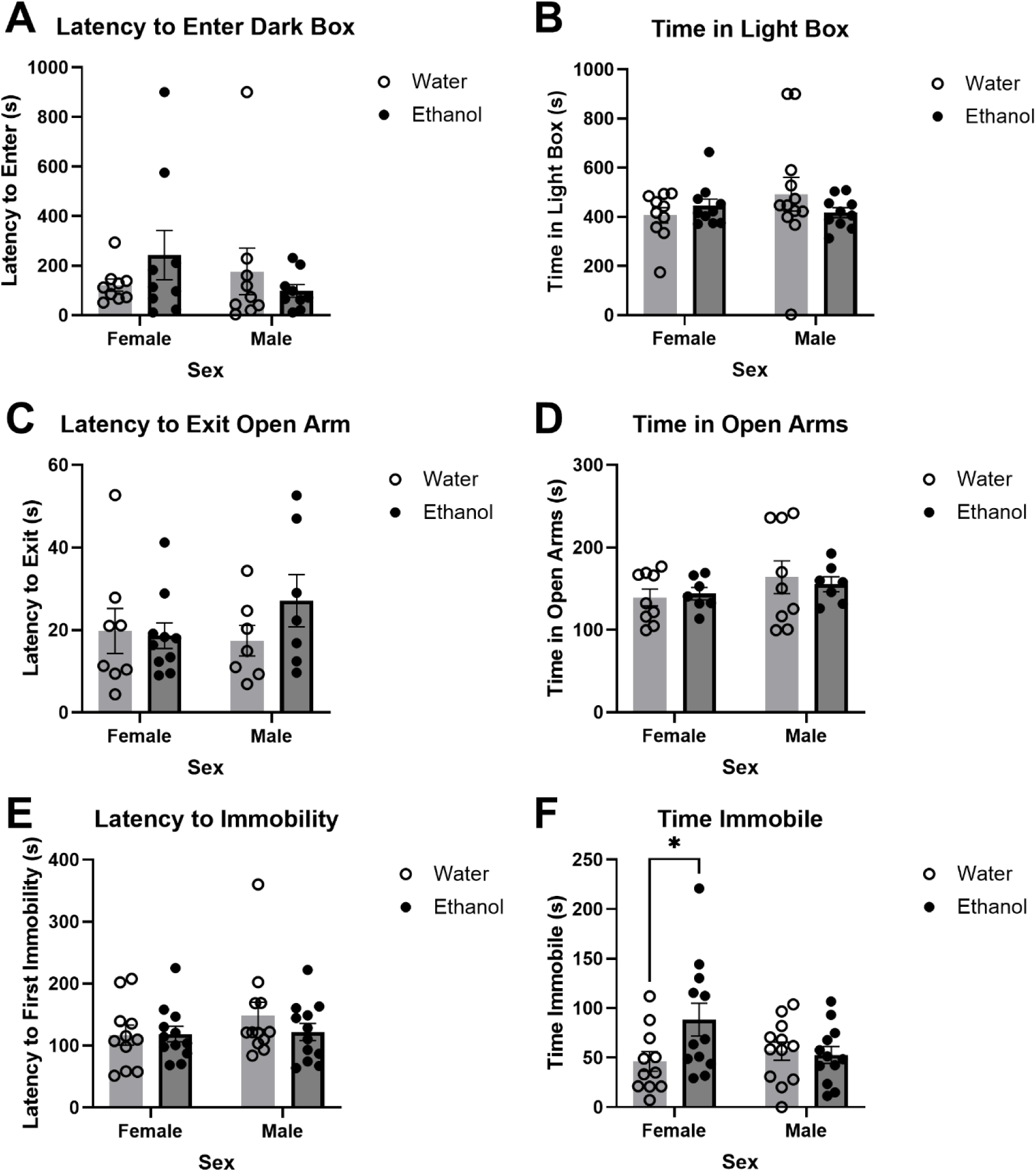
Negative affective-like behavior during protracted forced abstinence following adult-onset chronic ethanol consumption. A+B) No effects of ethanol abstinence or sex were observed in the light/dark box. C+D) No effects of ethanol abstinence were observed in the elevated plus maze. E+F) Forced swim test behavioral measures showed an increased time immobile for female ethanol mice compared to water controls, but no difference in males. *p<0.05.

In the forced swim test in adult-onset animals, there were no significant differences with latency to immobility (Fig. 3E). However, in examining the time immobile in the forced swim test, two-way ANOVA (Sex x Ethanol Group) indicated a trending interaction between Sex and Ethanol Group [F(1,43) = 3.948; p = 0.0533] (Fig. 3F). Fisher’s Least Significant Difference test indicates that ethanol group females spent significantly more time immobile in the forced swim test compared to water group females (p = 0.0142) whereas males do not show differences between the ethanol and water groups. Examining the unpaired mean difference between ethanol and water mice using estimation statistics indicated that female ethanol mice spent 42.2s more immobile during the 6min forced swim test than female water mice [95%CI 11.0, 83.1].

#### Negative Affective-Like Behaviors Following Adolescent-Onset Ethanol

Two weeks following removal of the ethanol bottle, we tested for sex differences in ethanol abstinence-induced negative affect-like behaviors in adolescent-onset mice. We observed no differences in behaviors in the light/dark box or elevated plus maze. Two-way ANOVAs (Sex x Ethanol Group) indicated no significant differences in either latency to enter the dark box and time in the light box in the light/dark box (Fig. 4A,B) and no significant differences in either latency to exit the open arm and time in the open arms in the elevated plus maze (Fig. 4C,D). In forced swim test in the adolescent-onset animals, two-way ANOVA (Sex x Ethanol Group) indicated a trending main effect of Ethanol Group on latency to immobility [F(1,41) = 4.058; p = 0.0506] with no main effect of Sex or interaction between Sex and Ethanol Group (Fig. 4E). Fisher’s Least Significant Difference test did not indicate any significant individual group differences. In examining the time immobile in the forced swim test, two-way ANOVA (Sex x Ethanol Group) indicated a significant main effect of Ethanol Group [F(1,40) = 9.970; p = 0.0030] (Fig. 4F). Fisher’s Least Significant Difference test indicates that ethanol mice spent significantly more time immobile compared to water for both females (p = 0.0292) and males (p = 0.0334). Examining the unpaired mean difference between ethanol and water mice using estimation statistics indicated that female ethanol mice spent 37.1s more immobile during the 6min forced swim test than female water mice [95%CI 7.86, 65.1] and that male ethanol mice spent 36.3s more immobile than male water mice [95%CI 1.31, 66.6].

**Fig. 4.**
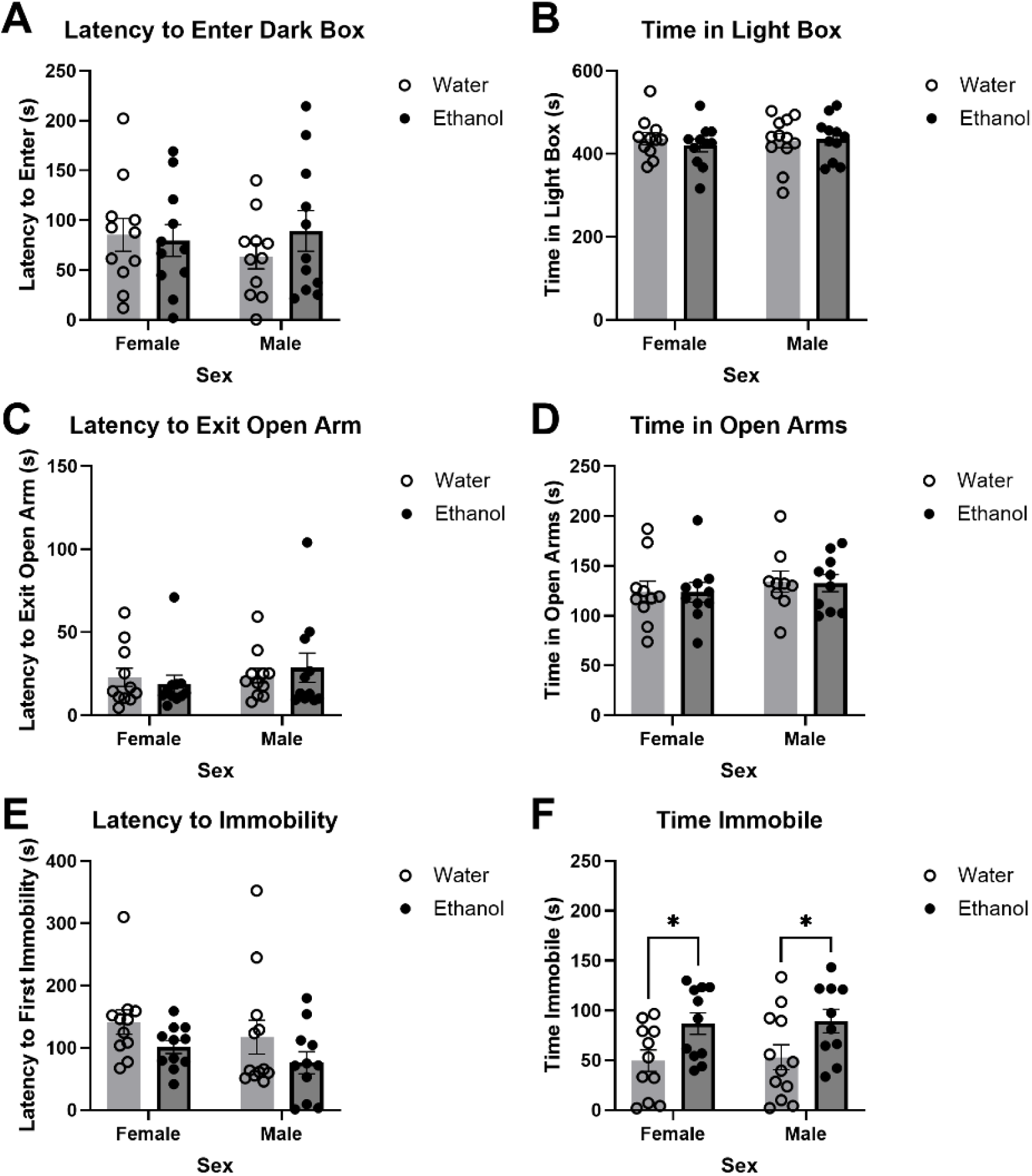
Negative affective-like behavior during protracted forced abstinence following adolescent-onset chronic ethanol consumption. A+B) No effects of ethanol abstinence or sex were observed in the light/dark box. C+D) No effects of ethanol abstinence were observed in the elevated plus maze. E+F) Forced swim test behavioral measures showed an increased time immobile for both male and female ethanol mice compared to water controls. *p<0.05.

## Discussion

This study tested the hypothesis that age of initiation of ethanol consumption (early adolescence vs. early adulthood) influences sex differences in voluntary ethanol intake/preference and protracted abstinence-induced negative affective behavior in mice. We predicted that starting to drink ethanol in early adolescence would normalize sex differences observed in mice that started drinking in adulthood. Indeed, our findings indicate that female mice that began drinking ethanol during adulthood consume significantly more ethanol and show a higher preference for ethanol over water (Fig. 1) as well as display higher abstinence-induced negative affective behavior (Fig. 3F) compared to male mice. However, when ethanol consumption began during early adolescence, both male and female mice exhibited similar levels of ethanol intake and preference (Fig. 2) as well as negative affective behavior during abstinence (Fig. 4). These results suggest that early adolescence may be a critical period in which ethanol consumption and the development of negative affect during abstinence is normalized between sexes, potentially due to developmental factors involved in equalizing vulnerability to the effects of ethanol.

Our results align with existing literature demonstrating higher ethanol consumption and preference in adult female mice compared to males in many voluntary consumption models males (Yoneyama et al. 2008; Centanni et al. 2018; Fulenwider et al. 2019; Dao et al. 2020; Bloch et al. 2020; Arnold et al. 2023; Perry et al. 2025; Yoon et al. 2025). This sex difference has been attributed to various factors, including hormonal influences and differences in ethanol metabolism. Our study provides novel insights by showing that these sex differences are not present when ethanol consumption begins during early adolescence. This finding suggests that the developmental stage at which ethanol consumption is initiated plays a crucial role in shaping future drinking behaviors and affective responses.

Differences in circulating sex hormone levels in adulthood may be related to the observed sex differences in ethanol consumption in adult animals. Estrogen has been linked to increased ethanol intake, with numerous studies supporting this connection (Lancaster et al. 1996; Ford et al. 2002; Hilderbrand and Lasek 2018; Satta et al. 2018; Bertholomey and Torregrossa 2019). Specifically, estrogen may act to enhance the rewarding effects of ethanol and plays a role in ethanol-induced activation of brain reward circuitry (Vandegrift et al. 2020). Relatedly, higher estrogen levels are associated with attentional bias toward ethanol cues (Griffith et al. 2023). Testosterone levels may also be involved in the observed sex differences in ethanol intake. Indeed, there is some evidence that testosterone suppresses ethanol intake (Vetter-O’Hagen et al. 2011; Vetter-O’Hagen and Spear 2011). Interestingly, ethanol consumption has been shown to increase estrogen levels in women, particularly following menopause (Handy et al. 2024) and increases aromatization of testosterone to estrogen in men (Emanuele and Emanuele 1998). Given that, in our study, adolescent female mice drank the same amount as adult female mice and the adolescent male mice increased their level of consumption to the level of the females (Fig. 2B), it is possible that these results may be explained by ethanol consumption shifting hormone levels towards higher estrogen/lower testosterone in the developing adolescents.

Adolescents across species are considered to have heightened reward sensitivity (Galván 2013). In general, adolescents have differential responses to dopamine (Tseng and O’Donnell 2006), higher dopamine receptor expression (Reynolds and Flores 2021) and enhanced excitability in dopamine neurons compared to adults (McCutcheon and Marinelli 2009). During adolescence, the brain is highly plastic and goes through major circuit reshaping (Fuhrmann et al. 2015; Dow-Edwards et al. 2019). As such, experiences during adolescence, including the experience of first ethanol reward, likely have a major impact on behavior later in life. It is possible that the adolescent-onset ethanol animals in our study learned to associate ethanol with reward early in their drinking, allowing them to have continued high ethanol preference and intake into adulthood (when sex differences would be present) at the later stages of the chronic drinking period.

Adolescents are also less sensitive to the aversive effects of ethanol (Spear and Varlinskaya 2010), including sedation (Draski et al. 2001), motor impairment (White et al. 2002), and conditioned taste aversion (CTA) (Anderson et al. 2010). These results are particularly interesting in the context of sex differences since adult male rats have heightened ethanol-induced CTA compared to females (Sherrill et al. 2011). These results may be related to ethanol clearance taking longer in adult males compared to both females and adolescent males (Varlinskaya and Spear 2004). However, if male rats are given ethanol during adolescence, they demonstrate reduced ethanol-induced CTA as adults (Sherrill et al. 2011). It is likely mice that began drinking in adolescence, particularly male mice, do not learn to associate ethanol with its aversive effects the way an adult male would, contributing to continued elevated ethanol consumption and preference.

Additionally, we observed increased negative affect-like behavior during forced abstinence in adult-onset ethanol female mice compared to adult-onset water controls, consistent with previous research, but did not observe differences in negative affect-like behavior in adult-onset ethanol males (Fig. 3). Notably, both male and female mice that began ethanol consumption during early adolescence displayed increased negative affect during forced abstinence (Fig. 4). These findings are particularly interesting in the context of clinical findings which list withdrawal and negative affect as two of the three leading causes for relapse in adolescents, along with social pressure (Cornelius et al. 2003). Negative affect during abstinence is a common phenotype following human adolescent ethanol intake (Winward et al. 2014). Consistent with our results, in a study of adolescents, both male and female students with a recent history of heavy drinking showed increased negative affect during abstinence, although males appeared to resolve negative affect more quickly (Bekman et al. 2013). Given these findings, it would be interesting to examine negative affect-like behavior in mice across multiple time points in abstinence to determine when these changes abate for both sexes.

While both male and female mice show similar abstinence-associated behavioral disturbances, it remains unclear whether these behaviors are driven by similar neural changes. Adolescent ethanol exposure produces a variety of sex differences in circuitry changes (Carzoli et al. 2019; Zhao et al. 2021). It is possible that the same behavioral outcome (immobility in the FST) in males and females could be driven by different neural mechanisms. Future research should examine changes in neural circuitry in abstinence following adolescent-onset ethanol.

One potential limitation of this study is that ethanol consumption data was measured manually by recording ethanol bottle weights every 48-72hrs. While we did not observe sex differences in adolescent-onset drinking behavior using these measurements, it is possible the sex differences do exist in patterns of drinking microstructure. It would be interesting to continuously examine drinking behavior using a home cage drinking monitor like the Lick Instance Quantifier Home Cage Device (LIQ HD) (Petersen et al. 2023) to better quantify subtle differences in ethanol drinking.

An additional limitation of this study is that all mice were singly housed to be able to measure ethanol consumption. Social isolation is a particularly strong stressor during adolescence (Almeida et al. 2021; Li et al. 2021) and the effects of social isolation may impact male and female animals in different ways. For example, adolescent social isolation has been shown to increase drinking in the dark during adulthood in males, but not females (Cullins and Chester 2024). Therefore, it is possible that the adolescent male mice in our study were more sensitive to the stress of social isolation compared to the female mice resulting in the observed elevation in ethanol consumption in the adolescent-onset males. Future research should examine adolescent drinking behaviors in a social setting now that it is possible to accurately quantify drinking in socially housed mice using newly developed technology, such as LIQ PARTI (Petersen et al. 2024).

The observed similarities in ethanol consumption, preference, and negative affective behavior between male and female mice initiated on ethanol during adolescence underscore the critical window of adolescence for the development of AUD. This period is marked by significant neurodevelopmental changes, which may render the brain more susceptible to the effects of ethanol. Our findings highlight the importance of early intervention and prevention strategies targeting adolescent ethanol use to mitigate the risk of developing AUD later in life, particularly in males.

Additionally, these findings enable the inclusion of both male and female rodents in research on voluntary ethanol consumption. This is crucial for the field, as it allows for a comprehensive evaluation of sex differences in mechanisms underlying ethanol consumption and the emergence of negative emotional states. Incorporating both sexes in this type of research has significant translational relevance, allowing for better understanding and the development of strategies to address ethanol-related issues across different populations.

Overall, our study emphasizes the critical importance of considering the timing of ethanol exposure and sex differences in understanding drinking behaviors and the development of negative affect during abstinence. Our findings highlight the pivotal role of adolescence as a critical window for the development of AUD, reinforcing the need for targeted strategies to address adolescent ethanol use. Continued exploration of the underlying mechanisms driving sex differences in ethanol consumption and negative affect, including hormonal influences and social and environmental factors, will enhance our understanding of AUD and lead to more effective approaches to treatment and prevention, ultimately improving public health outcomes.

